# DNA methylation patterns differ between free-living *Rhizobium leguminosarum* RCAM1026 and bacteroids formed in symbiosis with pea (*Pisum sativum* L.)

**DOI:** 10.1101/2021.10.28.466258

**Authors:** Alexey Afonin, Gribchenko Emma, Evgeny Zorin, Anton Sulima, Vladimir Zhukov

## Abstract

*Rhizobium leguminosarum* (*Rl*) is a common name for several genospecies of rhizobia able to form nitrogen-fixing nodules on the roots of pea (*Pisum sativum* L.) and undergo terminal differentiation into a symbiotic form called bacteroids. In this work, we compared the genomes of the free-living and differentiated forms of the *Rl* strain RCAM1026 using Oxford Nanopore long reads. No significant genome rearrangements were observed, but the relative abundances of replicons were different between the cell states. GANTC, GGCGCC and GATC methylated motifs have been found in the genome, along with genes for methyltransferases with matching predicted targets. Methylation patterns for the GANTC and GATC motives differed significantly depending on the cell state, which indicates their possible connection to the regulation of symbiotic differentiation. The GGCGCC motif was completely methylated in both bacteria states, and, apparently, is a target for the modification-restriction system. Currently, the methylation patterns in symbiotic bacteria are not extensively studied, so a further investigation of the topic coupled with gene expression data is needed to elucidate the function of differential methylation in terminal differentiation of *R. leguminosarum* and other rhizobia.

## Introduction

DNA methylation is an important epigenetic regulatory mechanism in prokaryotes. In addition to protecting the bacterial cell from phages and limiting horizontal transfer by digesting foreign DNA via restriction-modification (R-M) systems (Bickle & Krüger, 1993), methylation plays a significant role in the control of DNA replication and reparation, the cell cycle, and the adaptation due to the so-called phase variations (different methylation patterns in the bacterial population(Collier, 2009). For bacteria, DNA methylation is the primary means of epigenetic regulation (Beaulaurier et al., 2019; Casadesús & Low, 2006). Methylation systems typically consist of a DNA methylase and one or more DNA binding proteins that can overlap the target methylation site on DNA, subsequently blocking its methylation(Boye et al., 2000; Lobner-Olesen et al., 2003; Low & Casadesús, 2008).

Third generation sequencing is becoming more and more affordable, streamlining, among other things, the investigation of nucleic acid modifications such as methylation patterns. Currently, various aspects of methylation in bacteria are actively studied thanks to the development of technologies for high-precision, real-time sequencing of long molecules (McIntyre et al., 2019; Spadar et al., 2021), but there are still large knowledge gaps in this area. For instance, there is very little known about the significance of methylation in such an important process as the endosymbiosis of bacteria and plants.

Soil bacteria from the rhizobia group are able to partake in symbiosis with plants of the Fabaceae family, forming nitrogen-fixing root nodules (Downie, 2014). Inside these nodules, the bacteria differentiate into the symbiotic form called bacteroids, with the degree of differentiation depending on the plant-microsymbiont pair. In particular, members of the IRL (inverted repeat-lacking) clade of the *Papilionoidea* subfamily, such as pea, alfalfa, and clover, form the so-called indeterminate nodules, which are considered the most evolutionarily advanced (Coba de la Peña et al., 2018; Remigi et al., 2016; Sprent et al., 2013). Inside these nodules, bacteria undergo terminal differentiation, lose their ability to duplicate and become metabolically integrated with the host. Bacterial cells grow in size, increase their DNA content, assume irregular shapes and gain the ability to fix atmospheric N2, simultaneously losing the ability to return to a free-living state (Alunni & Gourion, 2016).

Not much is known about bacterial DNA methylation in symbiotic conditions. At the time of this writing, one publication and one preprint were available on the methylation patterns of rhizobia within a nitrogen-fixing nodule. In the article by Davis-Richardson et al., gene expression data was combined with the information about DNA methylation of *Bradyrhizobium diazoefficiens* strain USDA110 in symbiotic nodules of soybean (*Glycine max* (L.) Merr.) (Davis-Richardson et al., 2016). The results showed a specific methylation pattern associated with symbiotic conditions, but no further investigation was performed. The preprint by George C. diCenzo et al. describes the methylation patterns in multiple *Ensifer* strains in free-living and symbiotic conditions (George C. diCenzo et al., 2021). Both these works were using PacBio SMRT sequencing to investigate the DNA methylation. In our work, we applied Oxford Nanopore sequencing for genome assembly and methylation analysis of *Rhizobium leguminosarum (Rl)* strain RCAM1026 (A. Afonin et al., 2017), the effective symbiont of garden pea (*Pisum sativum* L.). This allowed us to discern between different types of nucleotides modifications, namely N6-methyladenine (6mA), N4-methylcytosine (4mC) and 5-methylcytosine (5mC), making it possible to obtain a more detailed picture of DNA methylation.

## Materials and Methods

### Bacteroid extraction

For the bacteroid isolation, plants of *P. sativum* cv. Frisson were inoculated with *R. leguminosarum* strain RCAM1026. After vegetation in quartz sand with mineral nutrition (A. M. Afonin, Leppyanen, et al., 2020) for 4 weeks, phenotypically mature (pink) nodules were collected in eppendorf tubes containing pre-chilled Tris-HCl/sucrose buffer (0.5 M sucrose-50 mM Tris-HCl [pH 8.0], dithiothreitol (10mM), and polyvinylpolypyrrolidone (5%)). The nodules were ground using mortar and pestle; the entire bacteroid isolation procedure, with the exception of the final stage, was carried out in sucrose buffer, as in (Catalano et al., 2004). To remove the plant cells debris, the crushed nodule material was filtered through a Miracloth-like material, followed by washing in the same buffer. The resulting suspension was centrifuged for 1 minute at 10,000 × g at +4°C. The symbiosome-containing pellet was resuspended in Tris-HCl/sucrose buffer. The resulting suspension was distributed into several Eppendorf tubes containing a “sucrose cushion” consisting of 1.5 M sucrose and 50 mM Tris-HCl [pH 8.0], and centrifuged for 30 sec. at 5,000 × g, +4°C. The upper phases, enriched with symbiosomes, were transferred into one polypropylene tube and centrifuged for 90 seconds at 10,000 g, +4°C, which made it possible to concentrate the symbiosome fraction into a pellet. The pellet was then resuspended in Tris-HCl/sucrose buffer and again applied to the “sucrose cushion” followed by centrifugation for 5 minutes at 10,000 × g, resulting in precipitation symbiosomes with bacteroids. To remove the peribacteroid membrane and isolate the bacteroids, the pellet was resuspended and centrifuged in a buffer containing Tris-HCl, but without sucrose. The resulting precipitate was frozen in liquid nitrogen and stored at −80°C for subsequent DNA extraction.

### DNA Isolation, Library Preparation, and Sequencing

For DNA isolation from the cell culture, one colony of *Rl* RCAM1026 was grown on an orbital shaker in a 50 ml flask with 10 ml of TY media for 18 hours at 28°C, 200 rpm. The cell density, as measured on the NanoDrop OneC spectrophotometer (Thermo FS), was 0.4. The cell culture and the bacteroids from the cv. Frisson nodules (see above) were further used for DNA isolation.

DNA from bacteroid samples and from the cell culture was isolated using a modified phenol-chloroform method as described in (Ausubel, 2002).

To obtain non-methylated DNA, the following whole-genome DNA amplification procedure was performed. The DNA isolated from the cell culture was sheared using a 32 gauge hypodermic needle (Meso-Relle, Italy). Fragment length analysis by the TapeStation system 4150 (Agilent, USA) showed the fragment distribution to be centered at around 13000 b.p. 30 ng of the sheared DNA were used for end-prep reaction using theNEBNext® Ultra™ II End Repair/dA-Tailing (E7546, New England Biolabs, USA). A double strand DNA fragment was formed by heating two oligonucleotides (NP_adapt_2_fw 5’ AAAGACAACCACGACTATAACGT 3’ and NP_adapt_2_rv 5’ CGTTATAGTCGTGGTTGTCTTT 3’) to 95°C in a water bath and letting them cool down at room temperature. The fragment was ligated to the end-prepped DNA using the Blunt/TA ligase (M0367, New England Biolabs, USA) according to manufacturer’s instructions. The NP_adapt_2_fw primer was used to amplify the DNA for sequencing. 8 rounds of PCR amplification were sufficient to obtain 200 ng of WGA DNA. The PCR reaction product was cleaned using AMPureXP beads (Beckman Coulter, USA) and the resulting DNA which was used for sequencing. The resulting DNA fragments were considered non-methylated, and the obtained data was used as non-methylated control in subsequent analysis after additional filtering.

Long-read whole genome sequencing was performed using a MinION sequencer (Oxford Nanopore Technologies, UK) in the Core Center ‘Genomic Technologies, Proteomics and Cell Biology’ in All-Russia Research Institute for Agricultural Microbiology (ARRIAM). The SQK-LSK109 Ligation Sequencing Kit with the EXP-NBD104 Native Barcoding Expansion 1-12 kit (Oxford Nanopore Technologies, UK) were used to prepare the library according to manufacturer’s instructions, omitting the DNA shearing step.

The reads were basecalled and demultiplexed using the Guppy_basecaller (v. 5.0.5), the genomes were assembled with Flye (v.2.9) (Kolmogorov et al., 2019) and polished with Racon (v1.3)(Vaser et al., 2017), Medaka (v.1.4.3), and Pilon (v. 1.23) (Walker et al., 2014) as in (A. M. Afonin, Gribchenko, et al., 2020).

The genome comparison was performed using MUMMER (v. 4.0.0)(Marçais et al., 2018). Sniffles structural variation caller (v1.0.12a) was used for the finding of genome rearrangements (Sedlazeck et al., 2018).

Megalodon toolkit (v 2.3.4) was used for methylation calling and pattern finding in the genome. The res_dna_r941_min_modbases-all-context_v001.cfg configuration file was used to get the methylation status of adenine and cytosine. The data was then visualised using custom R scripts. Additional methylation calling and motif analysis was performed using the Nanodisco (v 1.0.3) pipeline (Tourancheau et al., 2021).

The restriction enzyme database REBASE (“REBASE. The Restriction Enzyme Database”) was used to search for potential methyltransferase genes in the genomes of *Rl* strain RCAM0126.

## Results

### Genome assembly and comparison

In this study, the genome of the *Rl* strain RCAM1026 was assembled *de novo* using the long reads. The Nanopore sequencing yielded 283 m.b.p. of reads with N50 = 30 784 for the cell culture, and 1.4 g.b.p with N50 = 5 756 for the bacteroids; the reads were used for genome assembly as described in the materials and methods section. The genomes were compared using the MUMMER dnadiff script. No large genome rearrangements between the genome assemblies of the bacteroids and the cell culture were found, the lack of rearrangements was additionally confirmed using the Sniffles algorithm.

The RCAM1026 genome was assembled into 5 circular replicons. The sequences were rotated so that the chromosome started with the Ori sequence, and the other replicons began with the *RepA* gene. The resulting assembly consists of one chromosome and 4 plasmids (Table 1).

**Table 1.**

| Chromosome name | Length | CDS | tRNA genes | GC composition | Coverage in bacteroids | Coverage in cell culture |
| --- | --- | --- | --- | --- | --- | --- |
| Chromosome | 4921456 | 4736 | 51 | 61.09 | 1 | 1 |
| pRL10 | 629474 | 575 | 2 | 60.61 | 0.91 | 0.58 |
| pRL11 | 655637 | 625 | - | 60.89 | 0.94 | 0.5 |
| pRL12 | 838366 | 756 | - | 60.40 | 0.97 | 0.48 |
| Symbiotic plasmid | 268924 | 263 | - | 58.12 | 0.59 | 0.6 |

The replicon coverage was calculated in relation to the chromosome for each condition.

Although there were no structural differences between the free-living and symbiotic genomes, there was a difference in coverage. The coverage diagram for the cell culture and the bacteroid conditions (Figure 1) show uniform coverage of all the replicons. Relative coverage, however, differed between the conditions (Table 1), with the symbiotic plasmid (carrying the *nod* and *nif* clusters) being under-represented in the cell culture and the chromosome being over-represented in the bacteroids.

**Figure 1.**
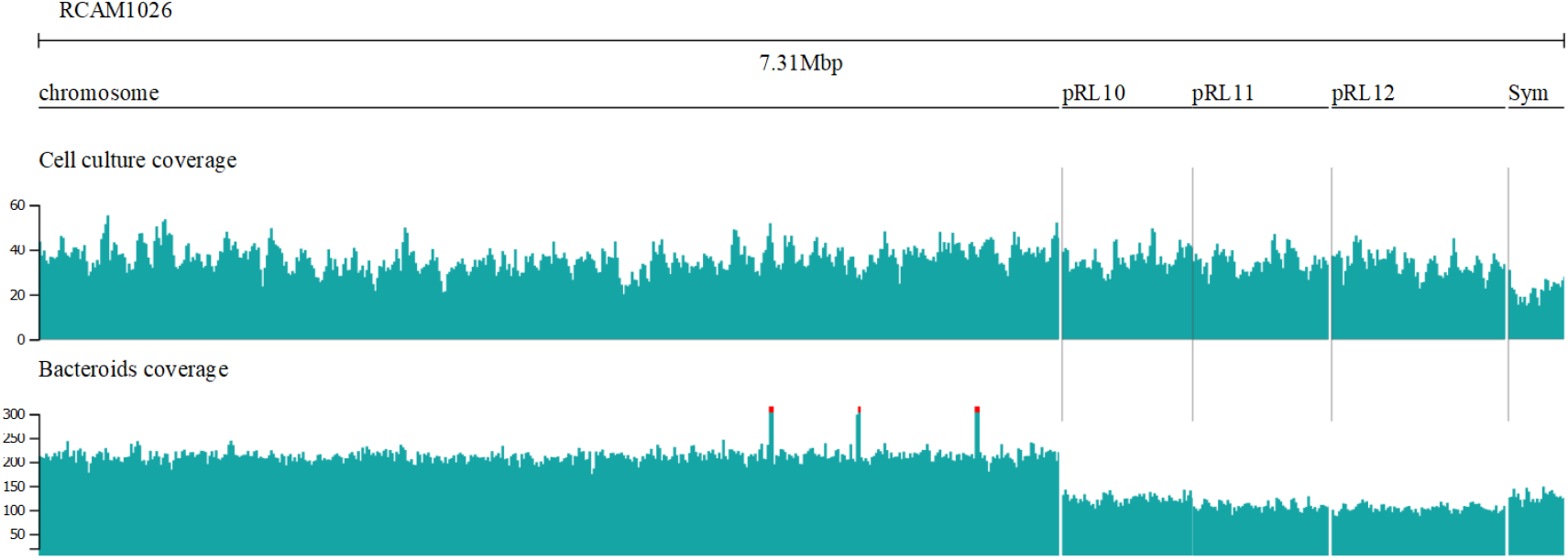
The coverage plot for the genome in two conditions. The plots represent the coverage statistics across the replicons.

### Methylation motives discovery

The cell culture genome assembly was used for this and subsequent steps. Methylation motives in the analysed samples were searched for using Nanodisco pipeline. GANTC, GATC, and GGCGCC methylated motifs were found to have a modified base. The precise mapping results are shown in Table 2. The GANTC motif showed a strong signal for 6mA base, which was in concordance with our expectation, as this type of methylation has been previously reported for the α-proteobacteria (Collier, 2009); however, the methylation of GGCGCC motif has been not reported for the *Rhizobuim leguminosarum* species complex.

**Table 2.**
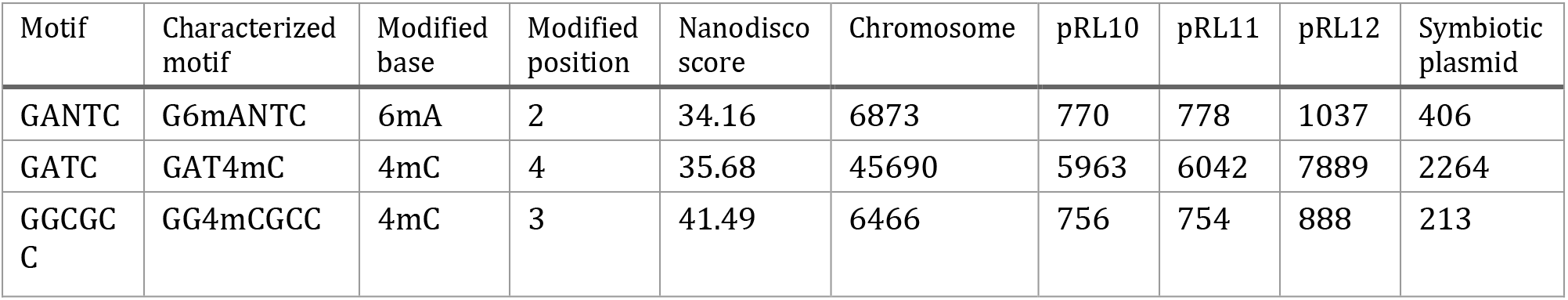
Motif distribution by replicon

### Genome methylation gene content

The annotated proteins in the RCAM1026 genome were BLAST-searched against the REBASE database. The gene most likely responsible for the methylation of GANTC motif is RCAM1026_000980 (97 % similarity to *M.retCII*). We found two copies of a methylase with predicted target motif GGCGCC (~52% similarity to *M.CcrNAIV*) - RCAM1026_001000 on a chromosome and RCAM1026_005751 on the pRL11 plasmid. In both locations the GGCGCC methylase gene is situated next to a Putative Type II restriction enzyme gene, suggesting the usage of the GGCGCC motif as a target for a restriction-modification system. Two copies of methyltransferase gene with predicted target motif GATC were found on the chromosome. The predicted modifications according to the database are G6mATC RCAM1026_002430 GAT5mC for RCAM1026_000801. All the found genes involved in methylation and their possible motifs are presented in Table 3. All the found genes are classified as Type II DNA methylation systems

**Table 3.**
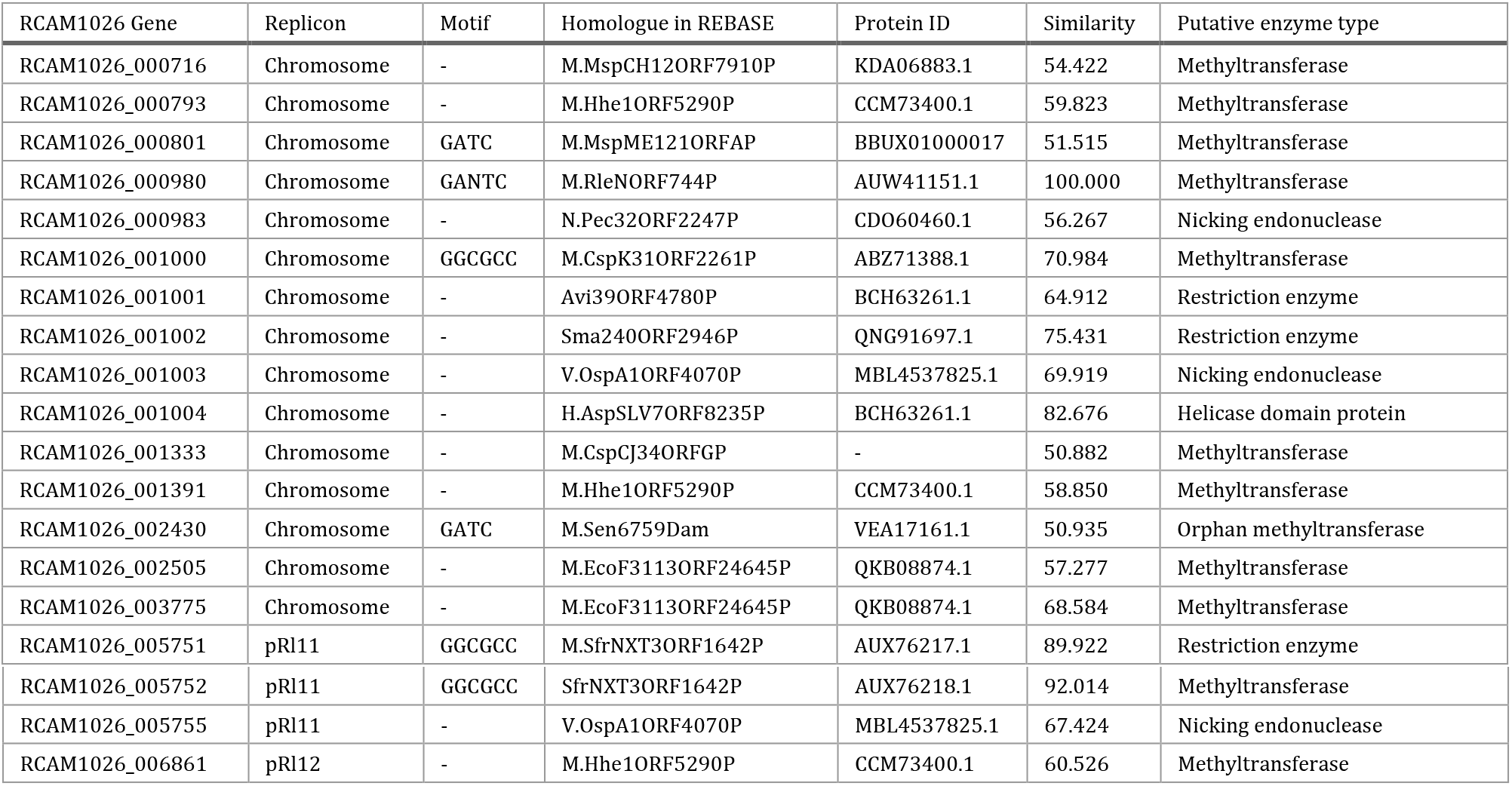
Genome methylation system content

### Methylation patterns

Since methylation can play a role in gene regulation, it was important to investigate whether the detected motifs are preferentially present in the promoter regions of the RCAM1026 genes. The results of the analysis for all three motifs are presented in Table 4.

**Table 4.**
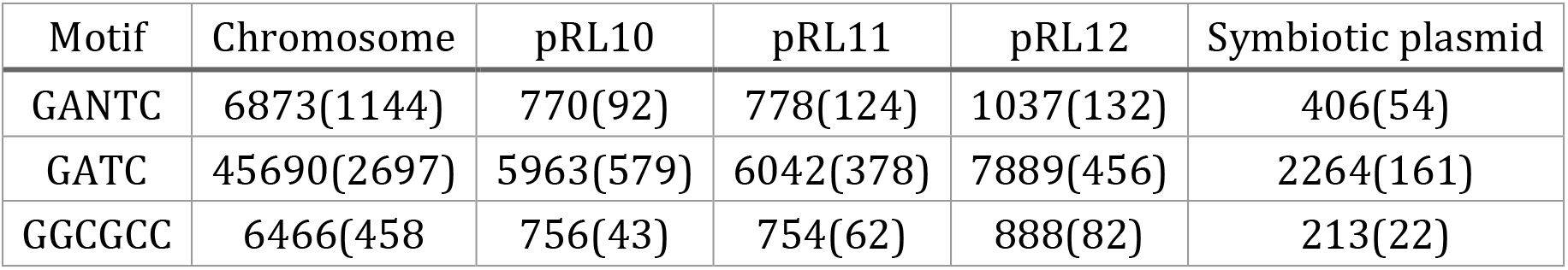
Methylation patterns intersection with genes and gene promoters

The number of motifs found in promoters is in parentheses.

Although the GANTC and the GGCGCC motifs are present in very similar numbers in the genomes, the GANTC motif is much more frequently found in the promoter regions. Genes with this motif in their promoter are likely to be regulated with GANTC methylation and play a role in the cell cycle.

### Genome-wide methylation pattern analysis

The precise methylation levels for each A (adenine) and C (cytosine) base as reported by the Megalodon pipeline were used for analysis of the DNA methylation. A model tuned for discerning A and C methylation simultaneously was used for methylation calling. As each read produced by the MinION sequencer corresponds directly to a fragment of native DNA, the data was not normalised. The levels of nucleotide methylation across the genome are presented in Figure 2.

**Figure 2.**
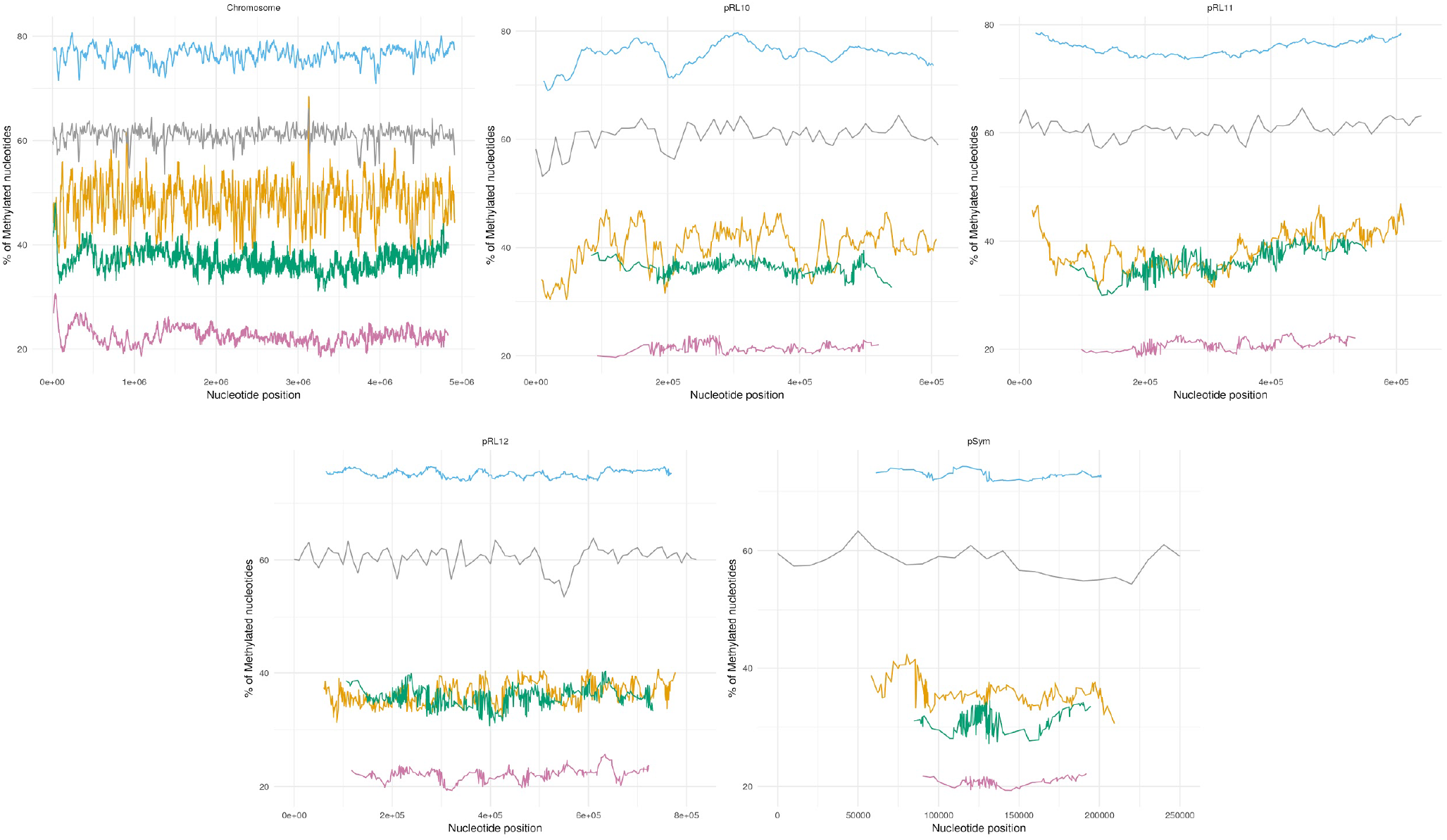
DNA methylation patterns. The plots represent the extent of methylation of adenine and cytosine the *R. leguminosarum* RCAM1026 genome shown using a 5 kb sliding window. The y scale shows the percentage of methylated reads mapped to the position. Blue - mA in bacteroids, purple - mA in cell culture, yellow - mC in bacteroids, green - mC in cell culture, grey - GC content.

The analysis shows that the levels of A methylation in the symbiotic state is much higher than in the free-living state, while the C methylation is much more similar between the two states.

### GGCGCC

The methylation patterns of the GGCGCC motif in two conditions are presented in figure 3. This motif is mostly methylated in all the replicons in both conditions, average cytosine methylation in this motif was observed to be 98%, compared to average 46% in bacteroids and 38 % in cell culture.

**Figure 3.**
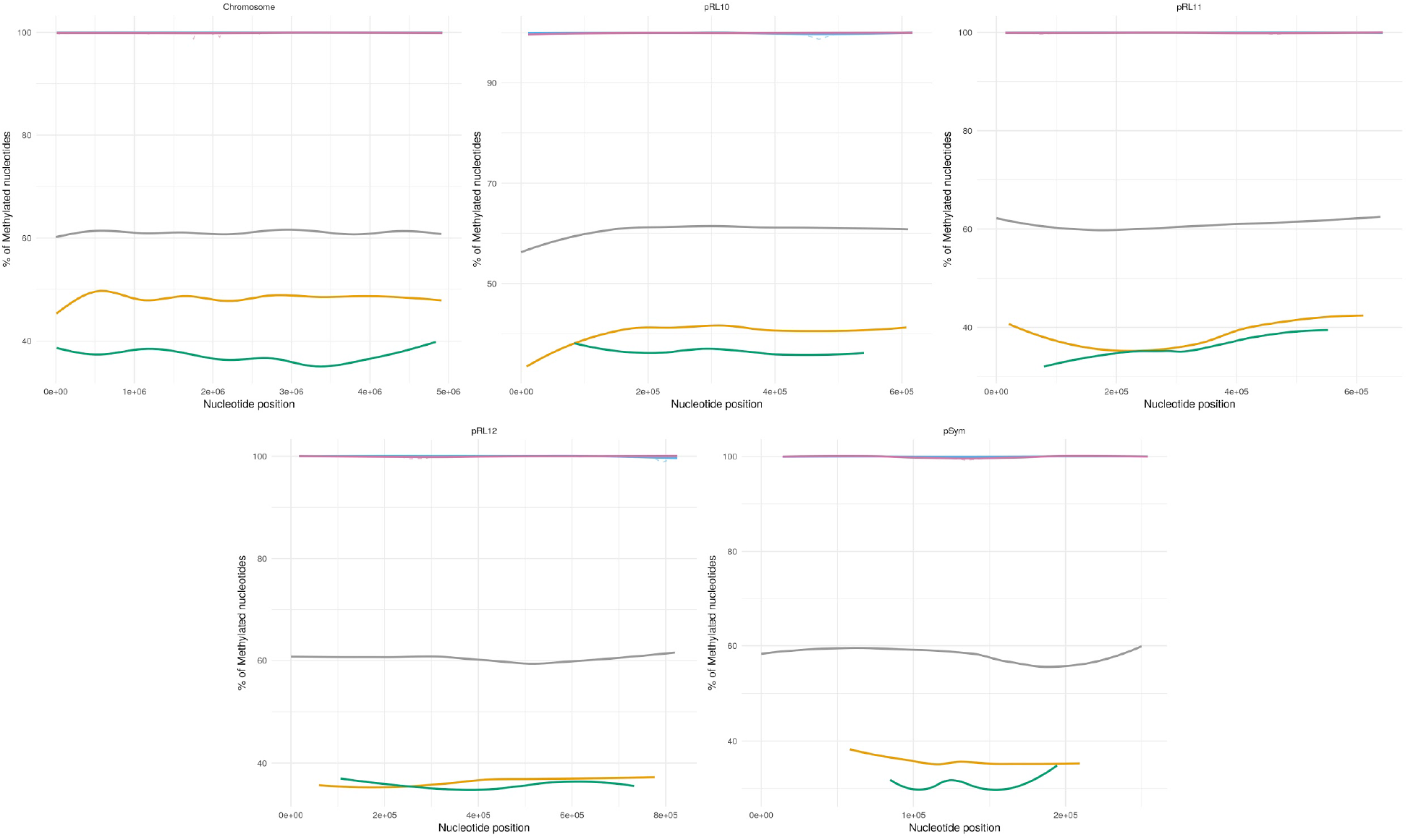
GGCGCC methylation across the replicons. Green - Blue *GGCGCC motif* methylation in bacteroids, purple - GGCGCC motif methylation in cell culture, yellow - methylation of cytosine in bacteroids, green - methylation of cytosine in bacteroids, grey - GC content. Solid lines - polynomial regression lines calculated using ggplot_smooth “loess” method, formula “y~poly(x,2)”, dotted lines - represent the extent of methylation using a 5 kb sliding window.

### GANTC

The methylation patterns of the GANTC motif in two conditions are presented in figure 4.

**Figure 4.**
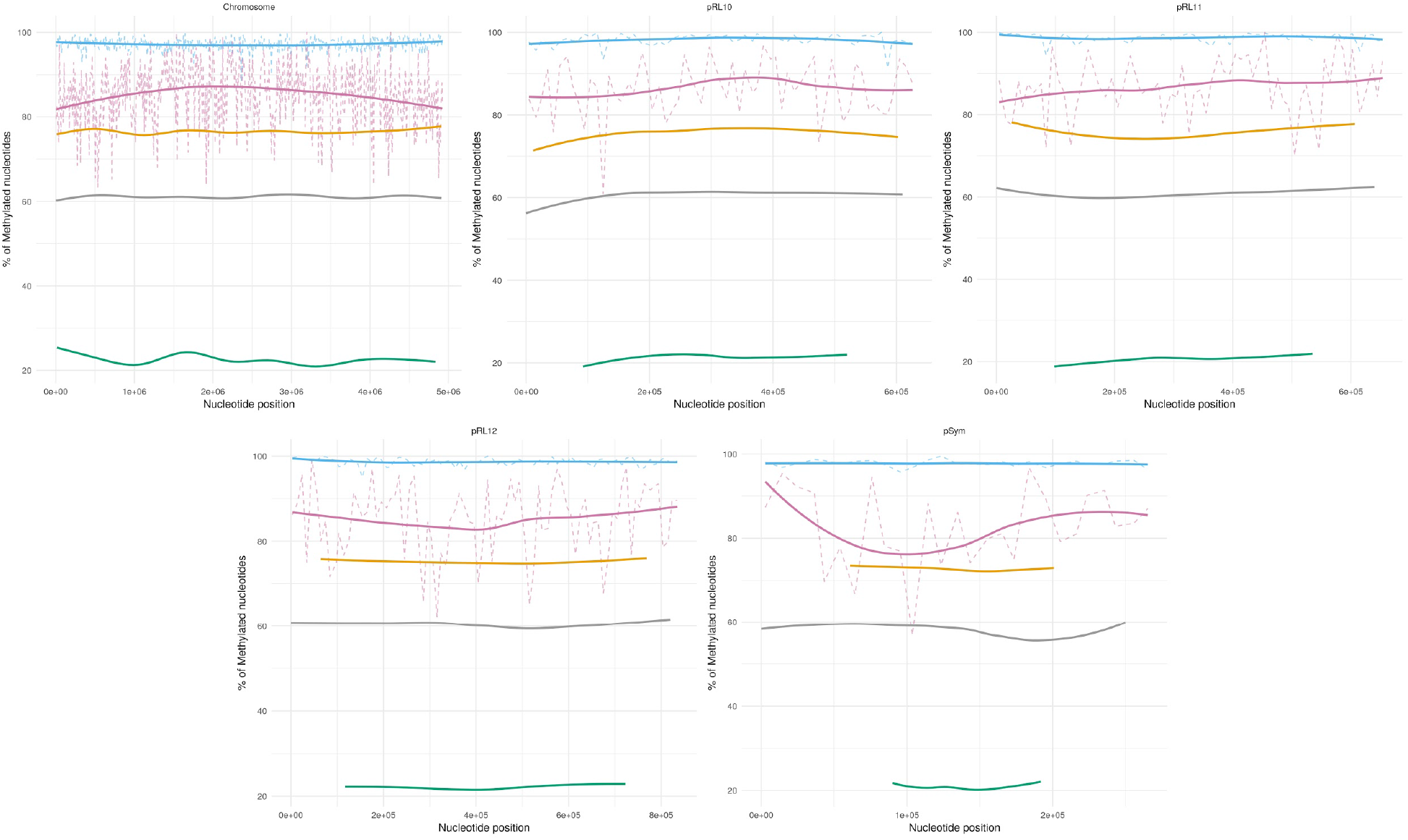
GANTC *methylation across the replicons*. Blue- GANTC motif methylation in bacteroids, purple - GANTC motif methylation in cell culture, yellow - methylation of adenine in bacteroids, green - methylation of adenine in cell culture, grey - GC content. Solid lines - polynomial regression lines calculated using ggplot_smooth “loess” method, formula “y~poly(x,2)”, dotted lines - represent the extent of methylation using a 5 kb sliding window.

In the bacteroids, the adenine methylation of the GANTC motif is at around 98% on all the chromosomes, higher than the 75% average for the A methylation, concordant with preferential methylation of A in this motif. For the cell culture, the GANTC motif shows increased methylation on the Ter region of the chromosome (the middle of the chromosome); no such pattern was observed for other replicons.

### GATC

The methylation patterns of the GATC motif in two conditions are presented in Figure 5.

**Figure 5.**
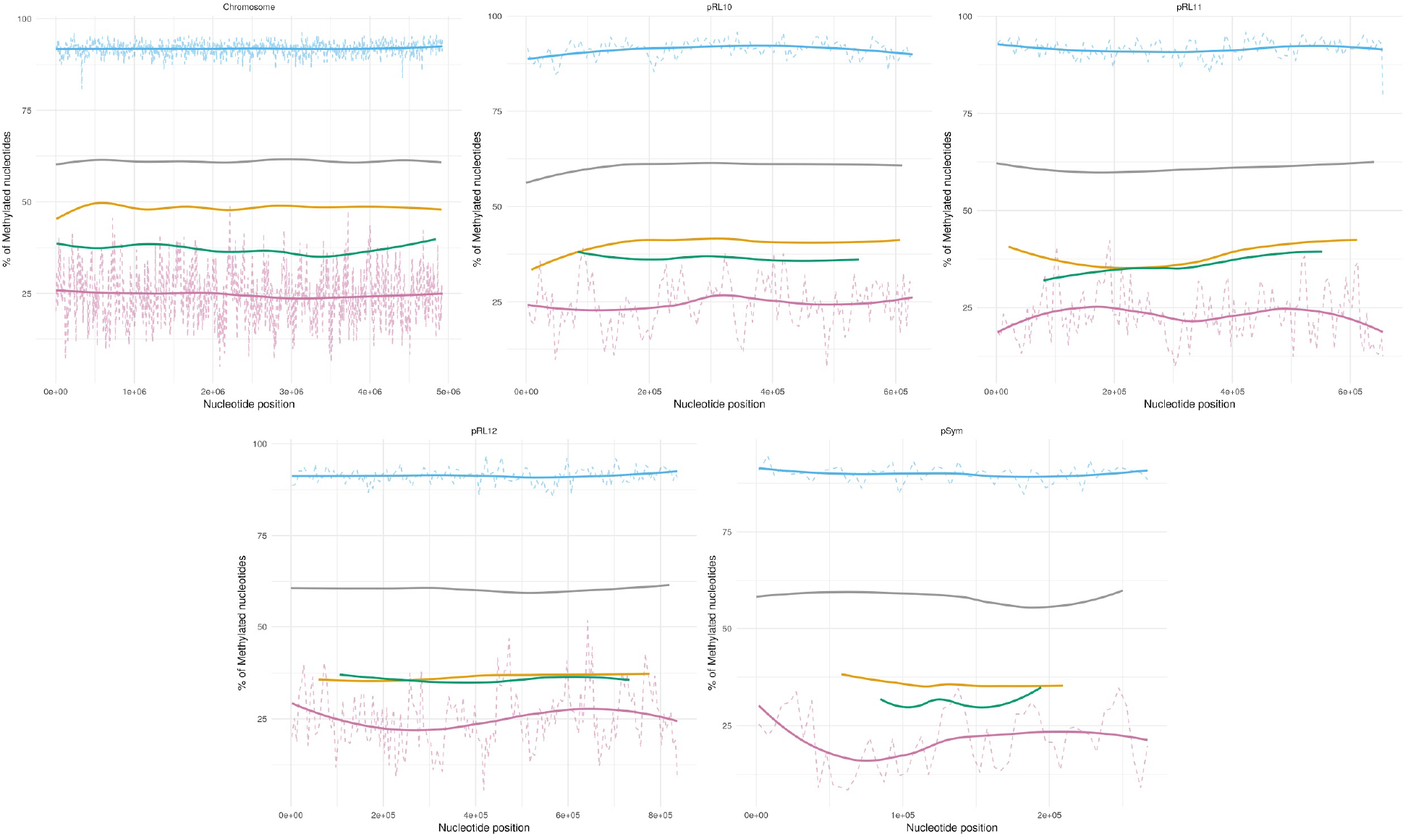
GATC methylation across the replicons. Green - GATC methylation in bacteroids, Violet - methylation in cell culture, red - methylation of cytosine in bacteroids, blue - methylation of cytosine in bacteroids, grey - GC content. Solid lines - polynomial regression lines calculated using ggplot_smooth “loess” method, formula “y~poly(x,2)”, dotted lines - the extent of methylation using a 5 kb sliding window.

The methylation patterns for the GATC motif are different from those of GANTC and GGCGCC motifs in both the investigated conditions. The methylation for this motif in bacteroids is around 80%, much higher than on average for all the replicons. For the cell culture, GATC motif is the only one with lower methylation than on average for the respective base.

## Discussion

Using Oxford Nanopore sequencing technology, the genome of the *Rhizobium leguminosarum* strain RCAM1026 was investigated in two different states - free-living cells and terminally differentiated bacteroids. The analysis did not show any genome rearrangements (in-dels, transpositions or loss of replicons) taking place in terminally differentiated cells. However, the genome coverage analysis showed significant differences in the relative abundances of replicons between the investigated states. The lower coverage of the symbiotic plasmid in free-living culture illustrates the relative ease with which the strain loses its symbiotic properties when propagated on solid media for a prolonged period of time. The observed phenotype may be a natural occurring population structure in which only a part of the population carries the plasmid, making the whole population more versatile in the environmental changes conditions (Heuer et al., 2008).

The equalisation of the plasmid content in the bacteroids is to be expected, as only the strain possessing the full array of symbiotically critical genes should be able to enter symbiotic state. The relatively higher coverage of the chromosome observed is similar to the observation made by George C. diCenzo et al. in the preprint (George C. diCenzo et al., 2021), where a similar pattern was described for the replicons in the bacteroids formed by *Ensifer* bacteria. Since the coverage may be influenced by a multitude of factors, the differences in DNA content in the bacteroids needs to be verified directly. The exact reason behind this abundance variation is unclear; one possible explanation is that the preferential reduplication of the chromosome makes it possible for the bacteria to increase the transcription levels of important metabolic genes, functionally doubling the number of gene copies in the cell.

Three motifs with significant methylation percentage were revealed: GANTC GGCGCC and GATC. For all the motifs, corresponding genes encoding methyltransferase enzymes were found in the genome. A few additional methyltransferase genes were also found, which is consistent with the overall methylation patterns for the adenine and cytosine not coinciding completely with the methylation patterns of the found motifs.

The Nanodisco pattern characterization showed GANTC to contain 6mA in the second position; GGCGCC and GATC contain 4mC base at third and fourth position, respectively. The methylation patterns in the two studied conditions are quite different. The 6mA is much more likely to occur in the bacteroids than in the cell culture, leading us to believe that 6mA is somehow involved in the terminal cell differentiation. Adenine-specific methylation has been well studied and shown to have diverse cellular roles (Ibryashkina et al., 2007; Wion & Casadesús, 2006). Indeed, the GANTC motif, known to be involved in the cell cycle regulation in proteobacteria (Collier, 2009), is fully methylated in the bacteroid genome which is consistent with the termination of cell division in the bacteroids. The bow-like pattern of GANTC methylation observed in the chromosome is in concordance with the fact that the methylation of DNA is dependent on the motif position in respect to replication origin (Ori site). No such pattern was observed for the plasmids.

The pattern of cytosine methylation is much more similar in the two conditions. Cytosine methylation seems to be less connected to the cell cycle and differentiation status. However, two found motifs with methylated cytosine behave very differently.

The GGCGCC pattern is most likely connected to the restriction-modification system (R-M). Taking into account the two annotated restriction-modification clusters, one on the chromosome and the other on the pRl11 plasmid, the system highlights the importance of this pattern for *R. leguminosarum*.

The presence of the R-M system on the pRL11 plasmid serves as a mechanism of plasmid persistence in bacterial cells (Naito et al., 1995). However, since the chromosome also has the R-M system with the same target motif, this mechanism would not work in this case. The plasmid R-M system may work in tandem with the chromosomal one, both participating in regulatory processes in the bacterial cell, possibly by enforcing a specific pattern of methylation (Vasu & Nagaraja, 2013).

Compared to the other two motifs, the GATC motif shows a unique pattern of methylation. The levels of methylation in cell culture are lower for this motif than for the cytosine on average, while in the bacteroids these levels are higher. This points at the possible activation of GATC-recognising methyltransferase in bacteroids, making GATC motif a prospective candidate for the terminal differentiation regulator.

Finally, the large number of putative methyltransferase genes with unknown target sequences are likely responsible for the methylation of bases not covered by the discovered motifs.

## Conclusions

Rapid development of single-molecule sequencing technologies opens up the prospects of deeper understanding of the genetic and epigenetic mechanisms that govern the living cells. In this work, we demonstrate that the differentiated bacteroids within the cells of pea symbiotic nodules have different methylation profiles compared to free-living bacterial culture, but do not have any DNA rearrangements such as deletions and/or duplications of parts of chromosome or plasmids. On the other hand, the overall coverage of replicons was not uniform, which possibly reflects the loss of the Sym plasmid in part of the bacterial population in free living culture, and higher levels of chromosome replication in bacteroids. The observed coverage differences for the symbiotic palsmid, previously also reported for the *Rl* strain A1 (A. M. Afonin, Gribchenko, et al., 2020), may speak in favour of the possible heterogeneity of bacterial population in nature being somehow beneficial.

Among the detected DNA methylation motifs, we found those present in both bacteroids and free-living cells. The discovery of corresponding DNA methyltransferase genes most likely targeting all found motifs in the RCAM 1026 genome indirectly supports our methylation pattern search and analysis. The GANTC motif was found in a large proportion of the gene promoters and was methylated in both the cell states, showing that *Rl* is not an exception from the previously described pattern. The existence of two restriction-modification system protein groups with target motif GGCGCC point at the importance of this system for the cell. The mechanism of the functioning of this R-M system is unclear.

GATC motif is the least studied of the three motifs found. Its exact function is unclear, but the preferential methylation of this motif in the bacteroid cells and very low methylation percentage in free-living conditions suggest some form of state-dependent activation mechanism of the found GATC methylase.

Since DNA methylation serves as a mechanism for regulation of gene expression, we expect that combining our data with the comprehensive transcriptome analysis will definitely help understanding the molecular genetic and epigenetic bases of bacteroid differentiation in the cells of legumes’ nodules.

## Author Contributions

Conceptualization, A.A. and V.Z; methodology, A.A., E.G., E.Z.; formal analysis, A.A.; data curation, A.A.; writing—original draft preparation, A.A., A.S., E.G., E.Z.; writing—review and editing, A.A., V.Z.; supervision, V.Z.. All authors have read and agreed to the published version of the manuscript.

## Institutional Review Board Statement

Not applicable.

## Informed Consent Statement

Not applicable.

## Data Availability Statement

This Whole Genome Shotgun project has been deposited at GenBank under the accession CP084696-CP084700. The raw Illumina data are deposited under the accession SRR16242825., Nanopore data for the cell culture (Nanopore signal data) - SRR16229315 (SRR16229311), WGA - SRR16229312 (SRR16229309), bacteroids SRR16229313 (SRR16229310). The version used in this paper is the second version of the genome.

## Literature cited

Afonin, A. M., Gribchenko, E. S., Sulima, A. S., & Zhukov, V. A. (2020). Complete Genome Sequence of an Efficient Rhizobium leguminosarum bv. viciae Strain, A1. Microbiology Resource Announcements, 9(19). https://doi.org/10.1128/mra.00249-20

Afonin, A. M., Leppyanen, I. v., Kulaeva, O. A., Shtark, O. Y., Tikhonovich, I. A., Dolgikh, E. A., & Zhukov, V. A. (2020). A high coverage reference transcriptome assembly of pea (Pisum sativum L.) mycorrhizal roots. Vavilovskii Zhurnal Genetiki i Selektsii, 24(4), 331–339. https://doi.org/10.18699/VJ20.625

Afonin, A., Sulima, A., Zhernakov, A., & Zhukov, V. (2017). Draft genome of the strain RCAM1026 Rhizobium leguminosarum bv. viciae. Genomics Data, 11, 85–86. https://doi.org/10.1016/j.gdata.2016.12.003

Alunni, B., & Gourion, B. (2016). Terminal bacteroid differentiation in the legume–rhizobium symbiosis: nodule-specific cysteine-rich peptides and beyond. New Phytologist, 211(2). https://doi.org/10.1111/nph.14025

Ausubel, F. M. (2002). Short protocols in molecular biology: a compendium of methods from Current protocols in molecular biology/. http://www.sidalc.net/cgi-bin/wxis.exe/?IsisScript=QUV.xis&method=post&formato=2&cantidad=1&expresion=mfn=004128

Beaulaurier, J., Schadt, E. E., & Fang, G. (2019). Deciphering bacterial epigenomes using modern sequencing technologies. Nature Reviews Genetics, 20(3). https://doi.org/10.1038/s41576-018-0081-3

Bickle, T. A., & Krüger, D. H. (1993). Biology of DNA restriction. Microbiological Reviews, 57(2). https://doi.org/10.1128/mr.57.2.434-450.1993

Boye, E., Løbner-Olesen, A., & Skarstad, K. (2000). Limiting DNA replication to once and only once. EMBO Reports, 1(6). https://doi.org/10.1093/embo-reports/kvd116

Casadesús, J., & Low, D. (2006). Epigenetic Gene Regulation in the Bacterial World. Microbiology and Molecular Biology Reviews, 70(3). https://doi.org/10.1128/MMBR.00016-06

Catalano, C. M., Lane, W. S., & Sherrier, D. J. (2004). Biochemical characterization of symbiosome membrane proteins from Medicago truncatula root nodules. Electrophoresis, 25(3), 519–531. https://doi.org/10.1002/elps.200305711

Coba de la Peña, T., Fedorova, E., Pueyo, J. J., & Lucas, M. M. (2018). The Symbiosome: Legume and Rhizobia Co-evolution toward a Nitrogen-Fixing Organelle? Frontiers in Plant Science, 8.https://doi.org/10.3389/fpls.2017.02229

Collier, J. (2009). Epigenetic regulation of the bacterial cell cycle. Current Opinion in Microbiology, 12(6).https://doi.org/10.1016/j.mib.2009.08.005

Davis-Richardson, A. G., Russell, J. T., Dias, R., McKinlay, A. J., Canepa, R., Fagen, J. R., Rusoff, K. T., Drew, J. C., Kolaczkowski, B., Emerich, D. W., & Triplett, E. W. (2016). Integrating DNA Methylation and Gene Expression Data in the Development of the Soybean-Bradyrhizobium N2-Fixing Symbiosis. Frontiers in Microbiology, 7.https://doi.org/10.3389/fmicb.2016.00518

Downie, J. A. (2014). Legume nodulation. Current Biology, 24(5). https://doi.org/10.1016/j.cub.2014.01.028

George C. diCenzo, Lisa Cangiol, Quentin Nicoud, Janis H.T. Cheng, Matthew J. Blow, Nicole Shapiro, Tanja Woyke, Emanuele G Biondi, Benoît Alunni, Alessio Mengoni, & Peter Mergaert. (2021). DNA methylation in Ensifer species during free-living growth and during nitrogen-fixing symbiosis with Medicago spp. BioRxiv.

Heuer, H., Abdo, Z., & Smalla, K. (2008). Patchy distribution of flexible genetic elements in bacterial populations mediates robustness to environmental uncertainty. FEMS Microbiology Ecology, 65(3). https://doi.org/10.1111/j.1574-6941.2008.00539.x

Ibryashkina, E. M., Zakharova, M. v, Baskunov, V. B., Bogdanova, E. S., Nagornykh, M. O., Den’mukhamedov, M. M., Melnik, B. S., Kolinski, A., Gront, D., Feder, M., Solonin, A. S., & Bujnicki, J. M. (2007). Type II restriction endonuclease R.Eco29kI is a member of the GIY-YIG nuclease superfamily. BMC Structural Biology, 7(1). https://doi.org/10.1186/1472-6807-7-48

Kolmogorov, M., Yuan, J., Lin, Y., & Pevzner, P. A. (2019). Assembly of long, error-prone reads using repeat graphs. Nature Biotechnology, 37(5), 540–546. https://doi.org/10.1038/s41587-019-0072-8

Lobner-Olesen, A., Marinus, M. G., & Hansen, F. G. (2003). Role of SeqA and Dam in Escherichia coli gene expression: A global/microarray analysis. Proceedings of the National Academy of Sciences, 100(8).https://doi.org/10.1073/pnas.0538053100

Low, D. A., & Casadesús, J. (2008). Clocks and switches: bacterial gene regulation by DNA adenine methylation. Current Opinion in Microbiology, 11(2). https://doi.org/10.1016/j.mib.2008.02.012

Marçais, G., Delcher, A. L., Phillippy, A. M., Coston, R., Salzberg, S. L., & Zimin, A. (2018). MUMmer4: A fast and versatile genome alignment system. PLOS Computational Biology, 14(1), e1005944. https://doi.org/10.1371/journal.pcbi.1005944

McIntyre, A. B. R., Alexander, N., Grigorev, K., Bezdan, D., Sichtig, H., Chiu, C. Y., & Mason, C. E. (2019). Single-molecule sequencing detection of N6-methyladenine in microbial reference materials. Nature Communications, 10(1). https://doi.org/10.1038/s41467-019-08289-9

Naito, T., Kusano, K., & Kobayashi, I. (1995). Selfish Behavior of Restriction-Modification Systems. Science, 267(5199). https://doi.org/10.1126/science.7846533

REBASE. The Restriction Enzyme Database. (n.d.). In Available online: http://rebase.neb.com/rebase/rebase.htm. Retrieved August 16, 2021, from http://rebase.neb.com/rebase/rebase.htm

Remigi, P., Zhu, J., Young, J. P. W., & Masson-Boivin, C. (2016). Symbiosis within Symbiosis: Evolving Nitrogen-Fixing Legume Symbionts. Trends in Microbiology, 24(1). https://doi.org/10.1016/j.tim.2015.10.007

Sedlazeck, F. J., Rescheneder, P., Smolka, M., Fang, H., Nattestad, M., von Haeseler, A., & Schatz, M. C. (2018). Accurate detection of complex structural variations using singlemolecule sequencing. Nature Methods, 15(6). https://doi.org/10.1038/s41592-018-0001-7

Spadar, A., Perdigão, J., Phelan, J., Charleston, J., Modesto, A., Elias, R., de Sessions, P. F., Hibberd, M. L., Campino, S., Duarte, A., & Clark, T. G. (2021). Methylation analysis of Klebsiella pneumoniae from Portuguese hospitals. Scientific Reports, 11(1). https://doi.org/10.1038/s41598-021-85724-2

Sprent, J. I., Ardley, J. K., & James, E. K. (2013). From North to South: A latitudinal look at legume nodulation processes. South African Journal of Botany, 89.https://doi.org/10.1016/j.sajb.2013.06.011

Tourancheau, A., Mead, E. A., Zhang, X.-S., & Fang, G. (2021). Discovering multiple types of DNA methylation from bacteria and microbiome using nanopore sequencing. Nature Methods, 18(5). https://doi.org/10.1038/s41592-021-01109-3

Vaser, R., Sović, I., Nagarajan, N., & Šikić, M. (2017). Fast and accurate de novo genome assembly from long uncorrected reads. Genome Research, 27(5), 737–746. https://doi.org/10.1101/gr.214270.116

Vasu, K., & Nagaraja, V. (2013). Diverse Functions of Restriction-Modification Systems in Addition to Cellular Defense. Microbiology and Molecular Biology Reviews, 77(1). https://doi.org/10.1128/MMBR.00044-12

Walker, B. J., Abeel, T., Shea, T., Priest, M., Abouelliel, A., Sakthikumar, S., Cuomo, C. A., Zeng, Q., Wortman, J., Young, S. K., & Earl, A. M. (2014). Pilon: An integrated tool for comprehensive microbial variant detection and genome assembly improvement. PLoS ONE, 9(11). https://doi.org/10.1371/journal.pone.0112963

Wion, D., & Casadesús, J. (2006). N6-methyl-adenine: an epigenetic signal for DNA–protein interactions. Nature Reviews Microbiology, 4(3). https://doi.org/10.1038/nrmicro1350

